# Computational drug repurposing identifies *N*-acetylglucosamine as a potential therapeutic compound for CLN3 Batten disease

**DOI:** 10.64898/2026.05.12.724723

**Authors:** Elina Casoli, Amali S. Fernando, Juliana CS. Chaves, Rebecca L. Johnston, Destiny Aranovitch, Sueanne Chear, Anthony L. Cook, Alex W. Hewitt, Eske M. Derks, Anthony R. White, Zachary Gerring, Lotta E. Oikari

## Abstract

Batten disease, also known as neuronal ceroid lipofuscinoses, is one of the most common causes of childhood dementia. It is characterized by the accumulation of lipofuscin in lysosomes, leading to loss of brain cell function, onset of dementia-like symptoms, vision loss and seizures and has extremely limited treatment options. Here, we performed computational drug repurposing analysis to identify existing compounds that may target Batten disease risk genes. A total of 81 candidate compounds were identified, 6 of which were selected based on clinical tractability for downstream testing in Batten disease (CLN3) iPSC-derived models. After confirming disease phenotype and drug candidate safety, CLN3 brain cell cultures treated with and without drug candidates underwent bulk RNA-seq to identify drug responses. One of the candidate drugs *N*-acetylglucosamine (GlcNAc) significantly upregulated Batten disease risk gene *CLN5* expression and several other lysosomal markers within CLN3 brain cells, and modulated several pathways implicated in lysosomal storage disorders. Importantly, GlcNAc significantly reduced lipofuscin burden in both CLN3 iPSC-derived neurons and astrocytes, supporting its investigation as an additional therapy for Batten disease.

## Introduction

Batten disease, also known as neuronal ceroid lipofuscinoses (NCL), is one of the most common causes of childhood dementia, characterized by a progressive deterioration of neurocognitive functions during childhood or adolescence, such as loss of cognition, memory, speech, and motor functions as well as behavioral and personality changes, seizures, disturbed sleep and loss of vision ^1,2^. In the U.S it is estimated that Batten disease affects approximately 2 – 4 out of every 100,000 births with around 14,000 children affected worldwide ^3^. Batten disease is caused by mutations in the ceroid lipofuscinoses neuronal (CLN) genes. There are 13 different CLN mutations described to date, each having distinct monogenetic defects in genes encoding proteins in the endolysosomal system ^4,5^. The mutations in CLN genes result in the toxic lysosomal accumulation of auto-fluorescent ceroid material (lipofuscin) containing both lipids and proteins in neurons and non-neuronal cells, leading to cell death ^6^. CLN3 Batten disease is the most common type of Batten disease, with an average age of 5 for onset of symptoms and a life expectancy of less than 30 years^2,4^. Most patients with CLN3 Batten disease carry a homozygous 1-Kb deletion in the CLN3 gene on chromosome 16, which interferes with the production of battenin (CLN3 protein), a transmembrane lysosomal protein involved in the trafficking of proteins and lipids as well as having a recently identified role in chloride efflux ^2,4,7^.

There is no cure for Batten disease and disease-modifying treatments are extremely limited. Only one approved disease-modifying therapy for CLN2 type Batten disease exists, cerliponase alfa (Brineura), which is an enzyme replacement therapy that slows disease progression ^8^. However, as the blood-brain barrier (BBB) poses a major challenge for delivering large molecular weight enzymes into the brain, cerliponase alfa is administered via invasive intracerebroventricular infusion directly into the brain ^9^. In addition to enzyme replacement therapy, current investigational therapeutic approaches across the childhood dementia spectrum include substrate reduction therapy, small molecule therapy, gene therapy and stem cell therapy ^1,10,11^.

Drug repurposing is an approach where new indications are identified for already existing, clinically approved drugs. As such, it circumvents the time-consuming and expensive process of developing a completely new drug. Large-scale omics data and powerful analysis tools available today accelerate drug repurposing analysis ^12-14^, making this approach a promising alternative in discovering potential new therapeutic compounds for the treatment of Batten disease. One approach to drug repurposing is the study of how known disease risk genes interact with other genes in a network, known as network propagation ^12,13^. The resulting networks can serve as a molecular substrate for the integration of drug-gene interaction data, to identify drug compounds that may perturb gene networks, rather than individual genes. Importantly, this approach can be tailored to identify compounds with favorable bioavailability metrics, such as molecular weight or lipophilicity, thereby maximizing the probability of discovering BBB permeable compounds.

Here we report a translational pipeline for identifying new potentially therapeutic compounds for Batten disease. We first performed computational drug repurposing and validated drug candidate effects in a CLN3 Batten disease induced pluripotent stem cell (iPSC)-derived brain cell model. Our results identified the dietary supplement N-acetylglucosamine (GlcNAc) as a potential new modulator of lysosome health with potential therapeutic implications in Batten disease or other lysosomal storage disorders.

## Results

### Computational drug repurposing analysis identifies new drug candidates for Batten disease

Computational drug repurposing analysis was performed to identify compounds that may target Batten disease causative genes and pathways (Fig. 1, Supplementary Table 1). Briefly, curated Batten disease risk genes were analyzed using brain-specific Bayesian gene co-expression networks, followed by functional annotation and drug–gene interaction mapping. The analysis resulted in the identification of 81 compounds (Supplementary Data 1), which consisted of modulators, inhibitors and activators of lysosomal genes. We further characterized the candidate compounds based on predicted BBB permeability and physicochemical properties, including Lipinski’s rule of 5 that identifies orally deliverable compounds based on the limits of four chemical properties ^15^.

**Figure 1.**
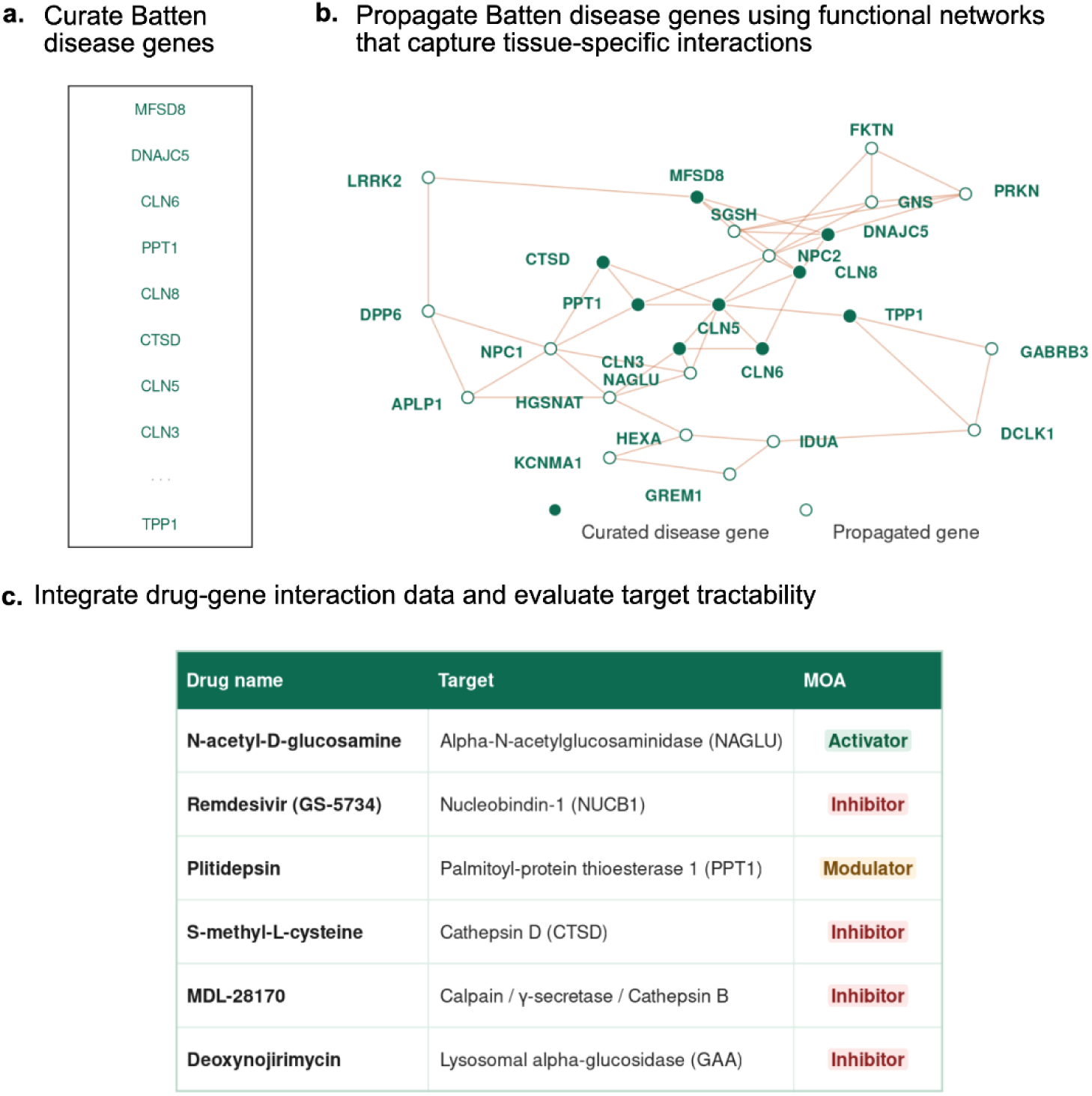
Overview of drug repurposing pipeline. **a)** Creation of an expanded Batten disease gene set using gene co-expression data from large-scale omics resources; **b)** Integration of drug-gene interaction data with expanded gene set; c) Drug candidate selection and categorization using experimental and clinical tractability information.

Six compounds targeting different lysosomal enzymes (including proteases, lipases and carbohydrases) were selected for downstream testing in a CLN3 iPSC-derived brain cell model (Table 1). *N*-acetylglucosamine (GlcNAc) is an activator of Alpha-N-acetylglucosaminidase (NAGLU), which plays a key role in degrading heparan sulfate and its dysfunction causes Mucopolysaccharidosis type IIIB ^16^. Remdesivir (Rem) is an inhibitor of nucleobindin-1 (NUCB1), with elevated NUCB1 associated with neuropsychiatric symptoms^17^. Plitidepsin (Pli) is a modulator of palmitoyl-protein thioesterase 1 (PPT1), a key lysosomal enzyme that breaks down fatty acids with defects in this enzyme causing CLN1 Batten disease ^18^. *S*-methyl-*L*-cysteine (SMLC) is an inhibitor of cathepsin D (lysosomal proteases) and a reported antioxidant that has shown therapeutic effects in a Parkinson’s disease model ^19^. MDL-28170 (MDL) is an inhibitor of proteases calpain, γ-secretase, and cathepsin B that has been reported to have anti-inflammatory and neuroprotective effects and has been tested previously in a CLN7 model ^20-22^. Deoxynojirimycin (Deoxy) is an inhibitor acid alpha-glucosidase (a carbohydrase) and its derivative miglustat is currently investigated for therapeutic effects in lysosomal storage disorders ^23,24^.

**Table 1.** Drug repurposing candidates selected for testing in Batten disease *in vitro* model.

| Drug name | Abbr. | Target | MOA | MW (Da) | Conc. used | Lipinski's rule of 5 violations |
| --- | --- | --- | --- | --- | --- | --- |
| **N-acetyl-D-glucosamine** | GlcNAc | Alpha-N-acetylglucosaminidase (NAGLU) | Activator | 221 | 1, 5 and 10 mM | 0 |
| **Remdesivir (GS-5734)** | Rem | Nucleobindin-1 (NUCB1) | Inhibitor | 603 | 1, 5 and 10 μM | 1 |
| **Plitidepsin** | Pli | Palmitoyl-protein thioesterase 1 (PPT1) | Modulator | 1110 | 1, 5 and 10 nM | 2 |
| **S-methyl-L-cysteine** | SMLC | Cathepsin D (CTSD) | Inhibitor | 135 | 1, 5 and 10<br>μM | 0 |
| **MDL-28170** | MDL | Calpain/γ-secretase/Cathepsin B | Inhibitor | 382.5 | 1, 5 and 10<br>μM | 0 |
| **Deoxynojirimycin** | Deoxy | Lysosomal alpha-glucosidase (GAA) | Inhibitor | 163.17 | 1, 5 and 10<br>μM | 0 |

Two of the drugs: Rem and Pli violated several of the Lipinski’s rule of 5 parameters, but were chosen for their previous clinical data. For example, Rem has been used as an anti-viral drug in the treatment of COVID-19 and it has been shown to have the ability to protect from neurological complications during viral infection ^25^. Pli has been mainly investigated as an anti-viral and an anti-cancer drug, it was chosen due to its predicted effects on the lysosomal pathway and the potential of using anti-cancer drugs as new drug repurposing options in neurodegenerative diseases ^26,27^. As such, these drugs are plausible candidates for correcting lysosomal dysfunction or having neuroprotective effects in Batten disease.

### CLN3 Batten disease *in vitro* model displays disease-associated phenotype

To generate a Batten disease-specific brain cell model for testing drug effects, iPSCs obtained from a person with CLN3 type Batten disease were used ^28^, along with an isogenic CLN3-corrected line ^28^ and an unrelated healthy control line HDFa ^29,30^. Neural progenitor cells (NPCs) were generated from each iPSC line and allowed to spontaneously differentiate to mixed cultures of neurons and astrocytes over a 30-day period to generate a bulk brain tissue model. Following differentiation, all three cell lines displayed expression of the neuron marker βIII-tubulin (TUBB3) and astrocyte marker glial fibrillary acidic protein (GFAP), along with the mature neuron marker microtubule associated protein-2 (MAP2) and mature astrocyte marker aquaporin-4 (AQP4) (Fig. 2a), indicating the presence of both neurons and astrocytes in the culture. Analysis of gene expression via qRT-PCR identified significantly higher expression of neuron markers doublecortin (*DCX*) (*P* < 0.01) and *TUBB3* (*P* < 0.001) in CLN3 cultures compared to CLN3-corrected and healthy cultures (Fig. 2b). Healthy cultures demonstrated significantly higher expression of astrocyte markers *GFAP* and S100 calcium binding protein B (*S100B*) (*P* < 0.001) compared to CLN3 and CLN3-corrected cultures (Fig. 2b). This likely reflects inherent differences between the iPSC lines in terms of their neuronal differentiation potential.

**Figure 2.**
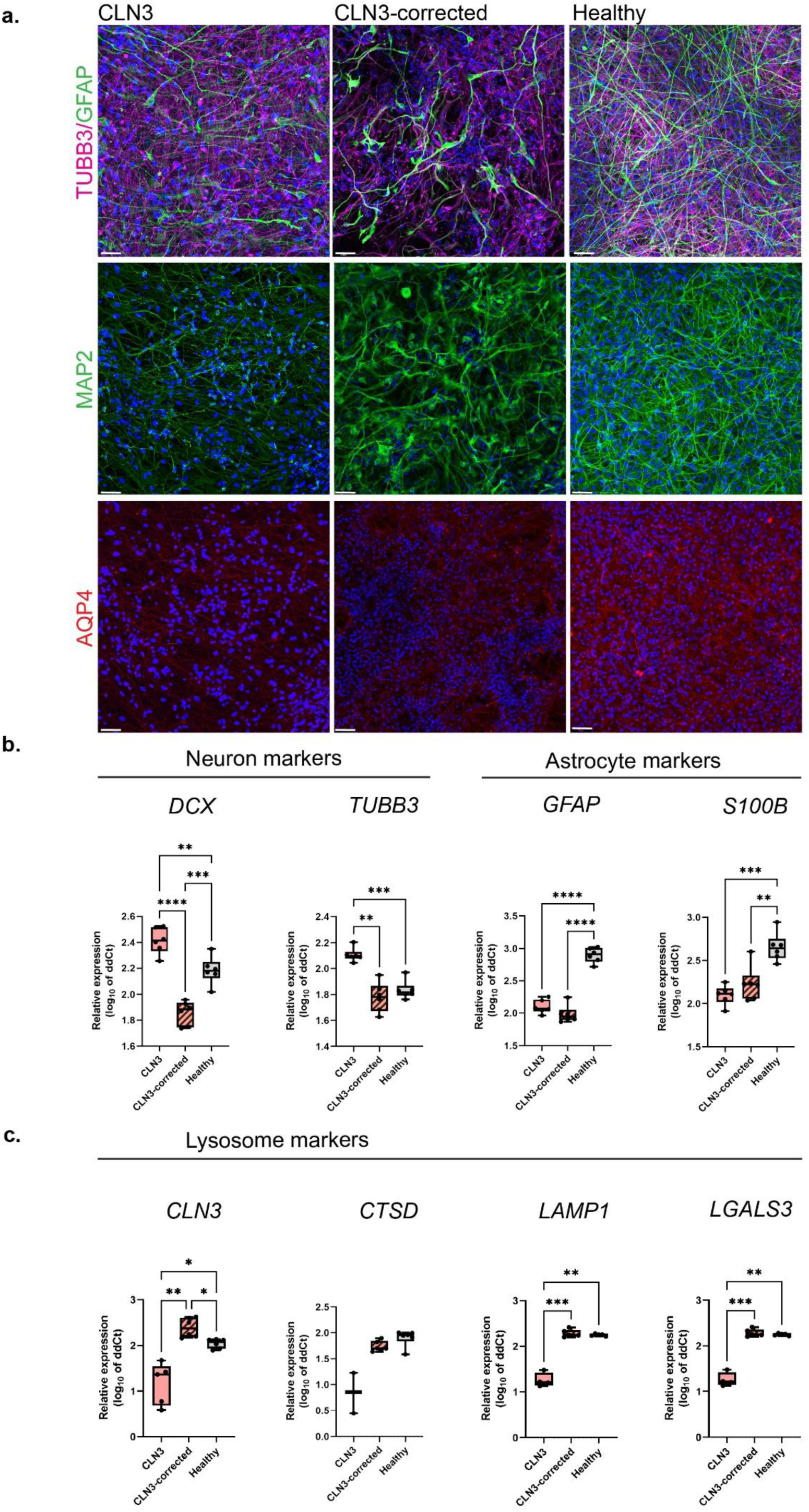
CLN3 iPSC-derived brain cell model characterization. **a)** Immunofluorescence of βIII-tubulin (TUBB3, magenta) and glial fibrillary acidic protein (GFAP, green), microtubule-associated protein 2 (MAP2, green) and aquaporin-4 (AQP4, red) in CLN3, CLN3-corrected and healthy iPSC-derived brain cell cultures (scale bar 50 µm). Relative gene expression of **b)** neuron and astrocyte markers doublecortin (*DCX*), *TUBB3*, *GFAP* and S100 calcium binding protein B (*S100B*) and **c)** lysosome markers CLN3 lysosomal/endosomal transmembrane protein (*CLN3*), cathepsin D (*CTSD*), lysosomal associated membrane protein (*LAMP1*) and galectin-3 (*LGALS3*) in CLN3, CLN3-corrected and healthy iPSC-derived brain cell cultures measured via qRT-PCR. Statistical analysis was performed using Welch ANOVA and defined as \**P* < 0.05, \*\**P* < 0.01, \*\*\**P* < 0.001 and \*\*\*\**P* < 0.001 (n = 6 independent differentiations per cell line).

Following neural marker analysis, the expression of Batten disease risk genes and lysosomal genes were compared between the cell cultures. Confirming disease phenotype, the *CLN3* gene was significantly downregulated in CLN3 cultures compared to CLN3-corrected and healthy cultures, with one of the CLN3 differentiation replicates expressing undetectable levels of *CLN3* (Fig. 2c). Cathepsin D (*CTSD*), a key lysosomal protease ^31^ also demonstrated reduced expression in the CLN3 cultures (non-significant), with only half of the studied samples demonstrating detectable expression levels (Fig. 2c). The lysosomal-associated membrane protein 1 (*LAMP1*), which plays a key role in regulating lysosome function ^32^ was significantly downregulated in CLN3 cultures compared to CLN3-corrected and healthy cultures, as was galectin-3 (*LGALS3*), which is key coordinator of lysosome repair and lysophagy ^33^ (Fig. 2c). These results indicate significantly impaired lysosomal function in CLN3 iPSC-derived brain cell cultures, also demonstrated in a previous study using the same iPSC line ^28^, supporting their suitability for therapeutic drug testing.

### Candidate drugs show minimal cytotoxic and neuroinflammatory effects

To ensure the safety of candidate drugs on brain cells, their effects on cell viability and inflammation were tested. For this, CLN3 brain cell cultures were exposed to drug treatments for 24 h after which supernatant was collected for LDH release assay, which measures the percentage of cell death. For each drug, three concentrations were tested based on ranges identified previously in the literature (Table 1). None of the vehicle or drug treatment conditions induced LDH release compared to untreated (UT) (Supplementary Fig. 1a) or compared to vehicle (both vehicle conditions pooled, given no statistically significant difference between the DMSO concentrations) (Fig. 3a).

**Figure 3.**
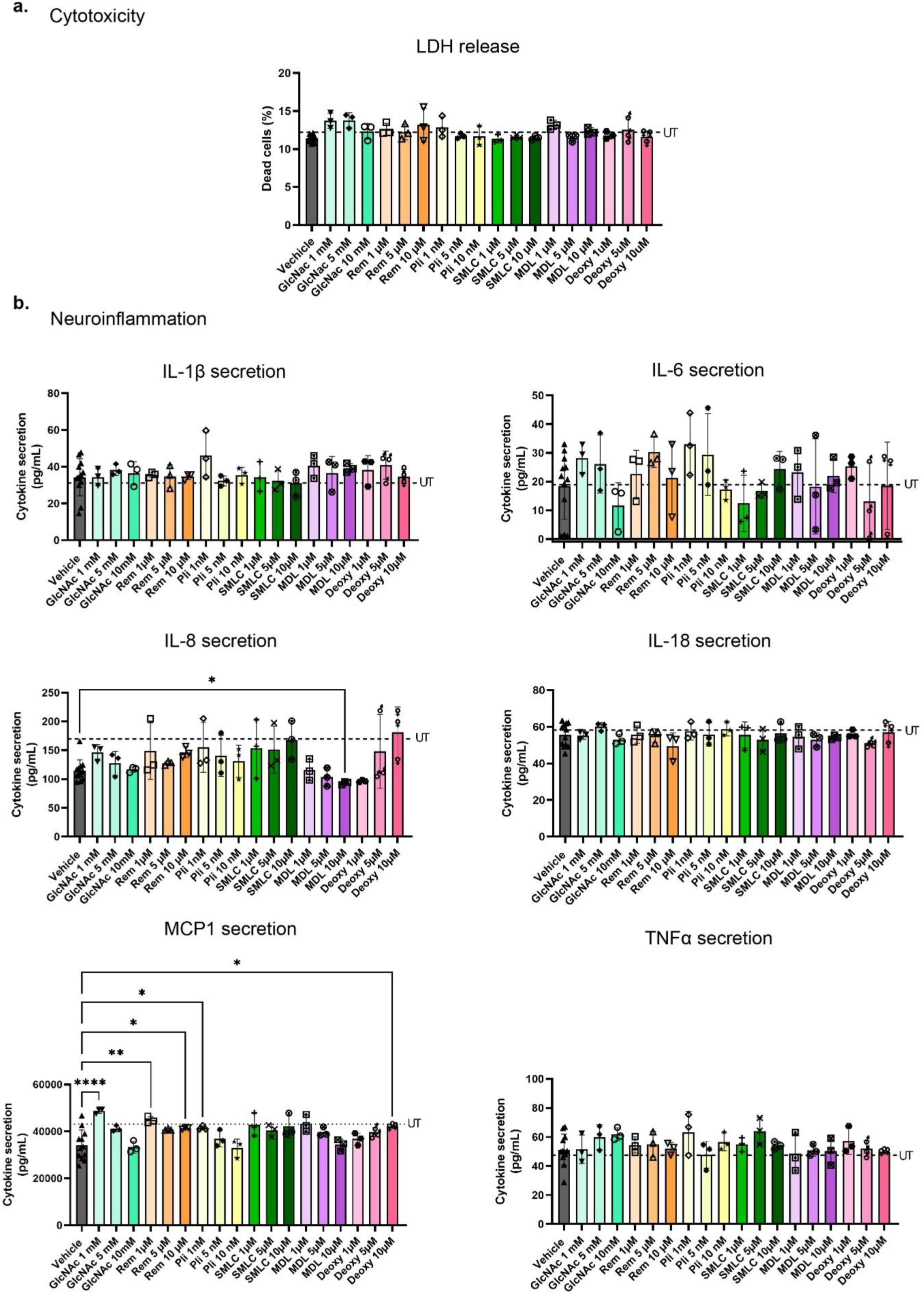
Effects of drug repurposing candidates on cytotoxicity and neuroinflammation. **a)** Lactate dehydrogenase (LDH) release cytotoxicity assay following treatment with drug repurposing candidates in CLN3 brain cell model. **b)** Cytokine secretion in CLN3 brain cell model following treatment with drug repurposing candidates. Statistical analysis was performed with one-way ANOVA and defined as \**P* < 0.05, \*\**P* < 0.01 and \*\*\*\**P* < 0.0001 (n = 3 independent replicates per treatment condition, high and low vehicle samples were pooled, comparisons only to vehicle shown with untreated condition (UT) shown as a dashed line).

Since altered neuroinflammation is characteristic of neurodegenerative diseases, the effect of candidate drugs on inflammatory ^34-36^ cytokine secretion was also investigated. None of the drugs significantly altered the secretion of key pro-inflammatory cytokines interleukin (IL)-1β, IL-6, IL-18 or tumor necrosis factor-α (TNFα) when compared to UT (Supplementary Fig. 1b) or vehicle (Fig. 3b). Interestingly, 10 µM MDL significantly (*P* < 0.05) reduced IL-8 secretion compared to vehicle, with 5 and 10 µM concentrations of MDL as well as 1 µm Deoxy significantly (*P* < 0.05) reducing IL-8 secretion compared to UT (Supplementary Fig. S2b). GlcNAc 1 mM, Rem 1 µM and 10 µM, as well as Pli 1 nM and Deoxy 10 µM were found to significantly increase MCP1 secretion compared to vehicle (Fig. 3b). Compared to UT, GlcNAc 1 mM also significantly elevated MCP1 secretion, with 1% DMSO and 10 mM GlcNAc both significantly reducing MCP1 secretion, suggesting the high DMSO concentration within these concentrations suppressed MCP1 secretion, as previously reported ^37^ (Supplementary Fig. 1b). These findings demonstrate that the drug concentrations used herein had minimal cytotoxic and inflammatory effects, with most effects associated with changes in IL-8 or MCP1 secretion.

### Differential expression analysis following candidate drug treatments

After confirming that none of the drugs induced cytotoxicity or marked neuroinflammation, CLN3 brain cell cultures treated with and without candidate drug compounds (GlcNAc 1mM, Rem 5 µM, Pli 10 nM, SMLC 10 µM, MDL 10 µM and Deoxy 10 µM) underwent bulk RNA-seq. Cells were exposed to drugs for 24 h after which RNA was extracted and processed for RNA-seq.

The number of differentially expressed genes (DEGs, FDR < 0.05) varied markedly across drug perturbation experiments in CLN3 brain cell cultures, ranging from relatively modest transcriptional changes for some compounds to broad transcriptome-wide perturbations for others. Rem (n = 3,293 DEGs) and GlcNAc (n = 1,846 DEGs) elicited the most amount of gene expression changes compared to vehicle, followed by Pli (n = 997 DEGs) and SMLC (n = 495 DEGs), with MDL (n = 17 DEGs) and Deoxy (n = 5 DEGs) treatment resulting in relatively few gene expression changes (Fig. 4a, Supplementary Data 2 - 7). We then investigated changes in the expression of Batten disease risk genes (*CLN1* to *CLN8* and *CLN10* to *CLN14*). Most notably, GlcNAc treatment resulted in a significant increase in *CLN5* expression (Fig. 4b). However, GlcNAc also downregulated *CLN4* (*DNAJC5*) and *CLN12* (*ATP13A2*) expression compared to vehicle, as did Rem, Pli, and SMLC treatment (Fig. 4b). Additionally, Pli treatment downregulated *CLN11* expression (*GRN*) (Fig. 4b), with no other significant effects observed for the other drugs.

**Figure 4.**
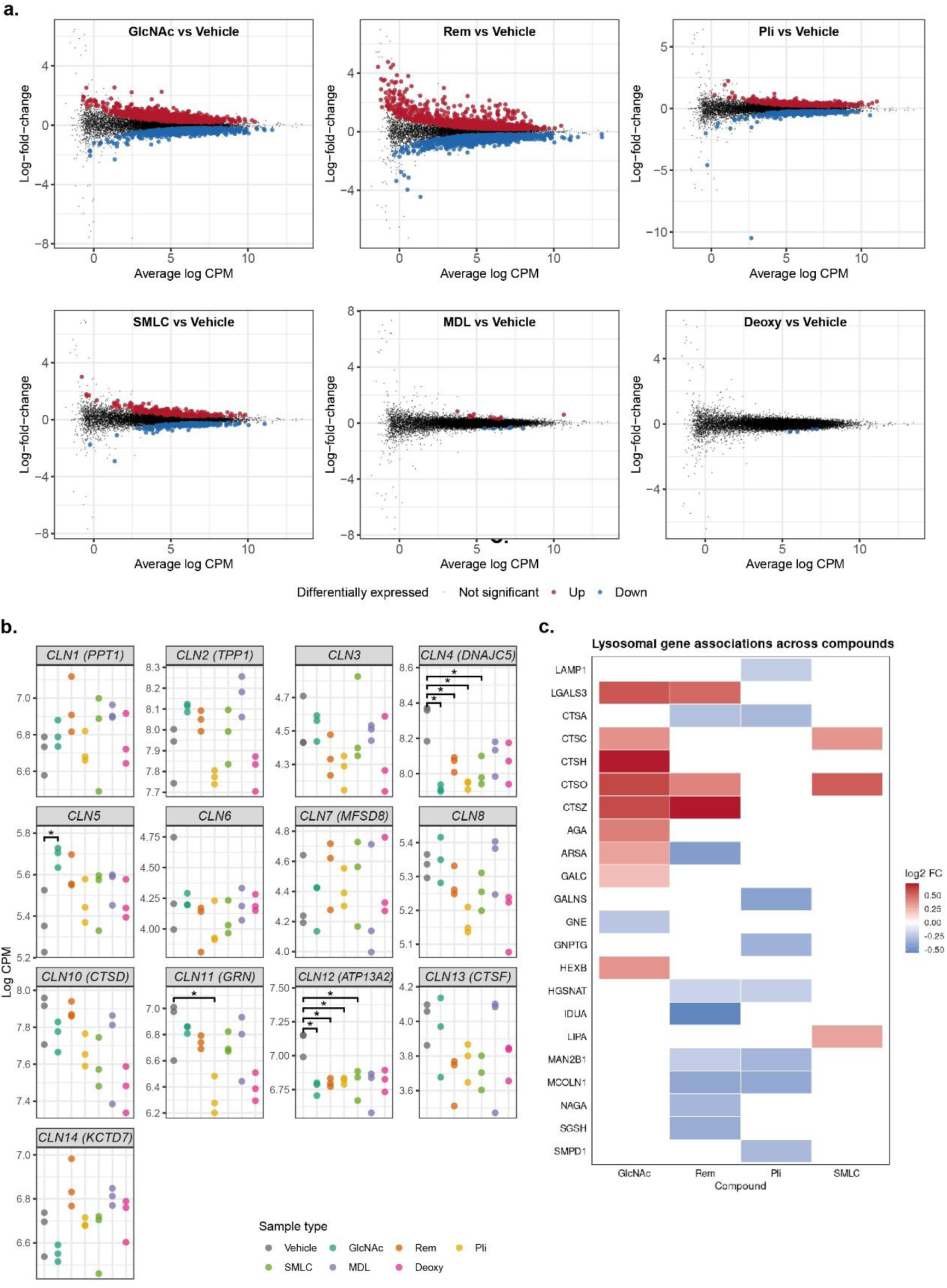
Differential expression analysis between candidate drug treatment versus vehicle control. **a)** Mean difference plots indicating differential expressed genes following each drug treatment (GlcNAc, Rem, Pli, SMLC, MDL and Deoxy) compared to vehicle. **b)** Scatter plots demonstrating expression of CLN genes across drug treatments compared to vehicle (* defined as FDR < 0.05) **c)** Heatmap of lysosomal and lysosomal storage disorder (LSD) associated genes (red = upregulated, FDR < 0.05, blue = downregulated, FDR<0.05, white = no significant fold-change, FDR > 0.05) in GlcNAc, Rem, Pli and SMLC treatment conditions compared to vehicle control. No significant changes in the selected genes were seen following MDL and Deoxy treatment.

We also investigated effects of the drug treatments on lysosome- and lipid-associated markers and enzymes, as well as genes associated with other lysosomal storage disorders (LSDs) previously listed ^38^. Interestingly, several of these markers were upregulated following GlcNAc treatment, including *LGALS3*, cathepsin (*CTS*) C, *CTSH*, *CTSO* and *CTSZ* (Fig. 4c). Importantly, GlcNAc significantly upregulated several genes associated with other LSDs, including *AGA* (aspartylglucosaminuria), *ARSA* (metachromatic leukodystrophy), *GALC* (Krabbe’s disease) and *HEXB* (Sandhoff disease), indicating lysosomal regulation (Fig. 4c) ^38^. In contrast, GlcNAc treatment significantly reduced *GNE* expression (Sialuria).

Rem treatment significantly upregulated *CTSO* and *CTSZ*, however, in contrast to GlcNAc, treatment Rem largely downregulated lysosomal and LSD-associated genes. These included *CTSA*, *ARSA*, *HGSNAT* (Sanfilippo syndrome C), *IDUA* (Hurler syndrome, Scheie syndrome), *MAN2B1* (Alpha mannosidosis), *MCOLN1* (Mucolipidosis IV), *NAGA* (Schinder’s disease) and *SGSH* (Sanfilippo syndrome A) (Fig. 4c). Pli treatment altered fewer lysosomal genes compared to GlcNAc and Rem with a downregulatory effect on the studied genes.

These included, *CTSA*, *GALNS* (Morquio syndrome A), *GNPTG* (Mucolipidosis III gamma), *HGSNAT*, *MAN2B1*, *MCOLN1* and *SMPD1* (Niemann-Pick disease A and B) (Fig. 4c). SMLC treatment significantly altered only three of the investigated genes, by upregulating *CTSC*, *CTSO* and *LIPA* (Lysosomal acid lipase deficiency) (Fig. 4c). No significant expression changes were identified on the investigated lysosomal genes following MDL or Deoxy treatment.

### Enriched pathway analysis following candidate drug treatments

Given most genes showed modest fold change values within each drug treatment, we next focussed on biological pathway-level results. Gene Set Enrichment Analysis (GSEA) was used to rank differential expression against Reactome pathways among the four compounds that elicited the most amount of gene expression changes (GlcNAc, Rem, Pli and SMLC). The top two positively and negatively enriched pathways (based on FDR < 0.05) from each treatment condition were identified.

GlcNAc treatment strongly induced positive enrichment in Extracellular matrix organization (R-HAS-1474244) and Collagen biosynthesis and modifying enzymes (R-HSA-1650814). Negative enrichment following GlcNAc treatment was observed neuronal and synaptic signalling pathways: Neuronal system (R-HSA-112316) and Transmission across chemical synapses (R-HSA-112310). (Supplementary Data 8), suggesting a shift toward a more structural or support-oriented cellular state (Fig 5a).

**Figure 5.**
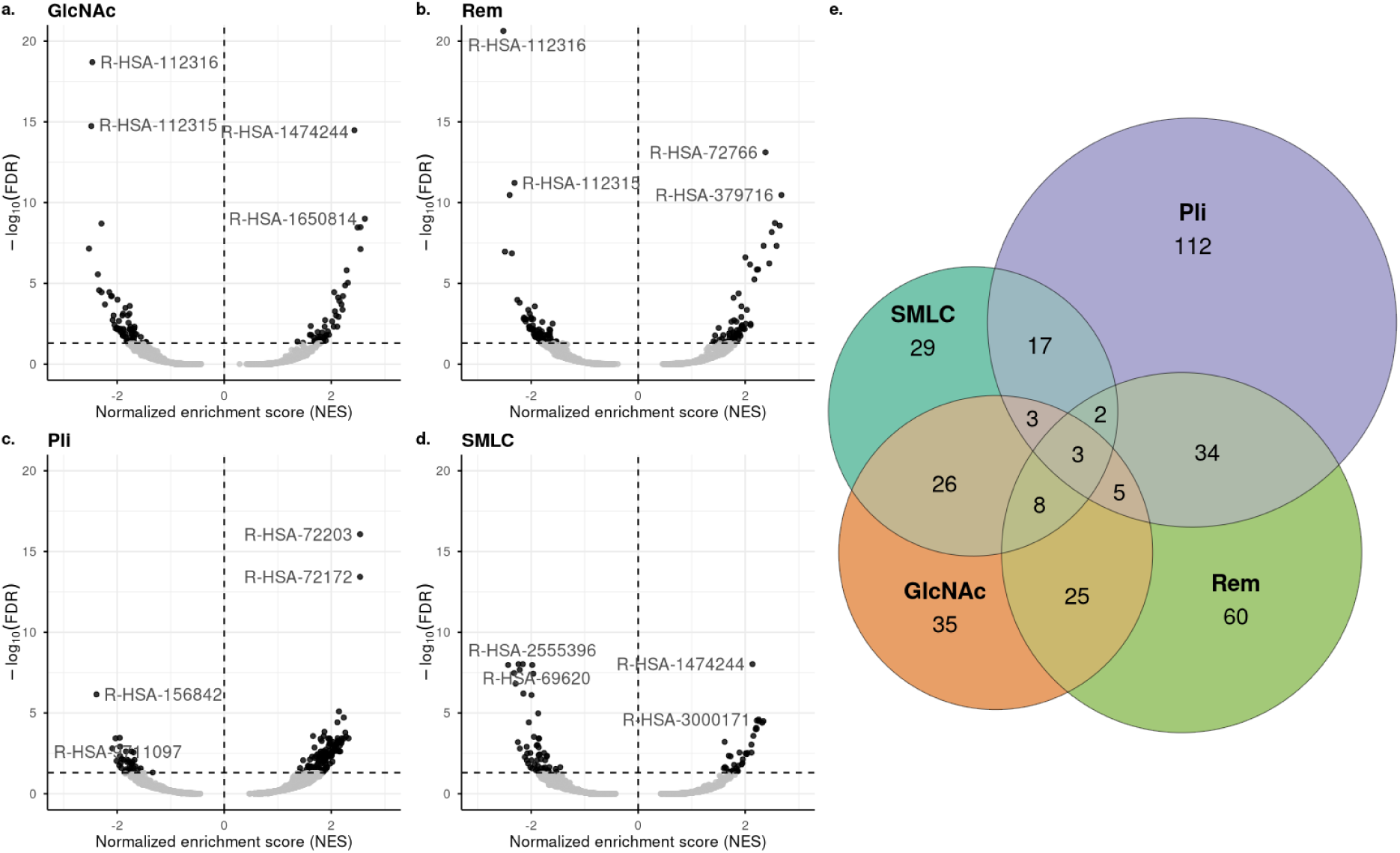
Gene-set enrichment analysis (GSEA) of differential gene expression for candidate drug treatments. Volcano plots of enriched pathways following a) GlcNAc, b) Rem, c) Pli and d) SMLC treatments, with the top 2 negatively or positively enriched pathways labelled by Reactome pathway identifiers. e) Venn diagram showing the number of shared and unique pathways following each drug treatment.

Rem treatment positively enriched Translation (R-HAS-72766) and Cytosolic tRNA aminoacylation (R-HSA-379716), and showed the same top negatively enriched pathways with GlcNAc (R-HSA-112316 and R-HAS-112315) (Supplementary Data 9) (Fig. 5b). Pli treatment positively enriched mRNA processing associated pathways, including: Processing of Capped Intron-Containing Pre-mRNA (R-HSA-72203) and mRNA splicing (R-HSA-72172) (Fig. 5c). Top negatively enriched pathways following Pli treatment were Eukaryotic Translation Elongation (R-HSA-156842) and Cellular response to starvation (R-HSA-9711097) (Supplementary Data 10) (Fig. 5c).

Similarly to GlcNAc, the top positively enriched pathway following treatment with SMLC was Extracellular matrix organization (R-HSA-1474244). In addition, the Non-integrin membrane - ECM - collagen pathway was positively enriched (R-HSA-3000171) (Fig. 5d). Top negatively enriched pathways following SMLC treatment were Mitotic metaphase and anaphase (R-HSA-2555396) and Cell cycle checkpoints (R-HSA-69620) suggesting effects on the cell cycle (Supplementary Data 11) (Fig. 5d).

We then assessed the number of unique and shared pathways between GlcNAc, Rem, Pli and SMLC treatments through a Venn diagram analysis. There were three shared pathways across all treatments (Fig. 5e), which were Neuronal System, Dopamine neurotransmitter release cycle and Serotonin neurotransmitter release cycle, all negatively enriched across the treatments (Supplementary Data 12). Rem and Pli treatments resulted in the highest numbers of shared pathways (n = 34), followed by GlcNAc and SMLC (n = 26), GlcNAc and Rem (n = 25) and Pli and SMLC (n = 17) (Fig. 5e). The highest number of unique pathways were identified in Pli treatment (n = 112), followed by Rem (n = 60), GlcNAc (n = 29) and SMLC (n = 29) treatments (Fig. 5e).

We then investigated unique pathways in each treatment condition with relevance to Batten disease or lysosomal storage disorders to gain insight into potential unique effects of each drug in CLN3 brain cells (summarized in Table 2). Interestingly, GlcNAc treatment positively enriched O-linked glycosylation, fatty acid metabolism associated pathways and keratan sulfate degradation pathways, which have direct relevance to lysosomal storage disorders and neurodegeneration (Table 2). Several key authophagy and mitophagy associated pathways were, however, negatively enriched, which may not be ideal for lysosomal function (Table 2). Rem treatment positively enriched protein processing pathways which might support protein breakdown (Table 2). However, several glycosaminoglycan degradation pathways, critical for reducing storage material build up in lysosomes, were negatively enriched (Table 2.). Pli positively enriched the Organelle biogenesis and maintenance pathway and negatively enriched Mucopolysaccharidoses, which could have potential therapeutic implications on lysosomal health (Table 2). However, Pli had several potentially undesired effects, including the positive enrichment phagosomal suppression and the negative enrichment in various transport, trafficking and lipid metabolism pathways linked to disease mechanisms of Batten disease and lysosomal storage disorders (Table 2). SMLC treatment resulted in the unique enrichment of only a small number of pathways with Batten disease or other lysosomal storage disorder relevance, including positive enrichment in cholesterol biosynthesis and steroid metabolism, and negative enrichment in intracellular trafficking (Table 2).

**Table 2.** Unique pathways following candidate drug treatment with relevance to Batten disease and other lysosomal storage disorders.

|  | *Reactome ID* | *Pathway name* | *Padj.* | *NES* | *Ref* |
| --- | --- | --- | --- | --- | --- |
| *GlcNAc* | R-HSA-5173105 | O linked glycosylation | 0.010 | 1.74 | 39,40 |
| *GlcNAc* | R-HSA-77288 | Mitochondrial fatty acid beta oxidation of unsaturated fatty acids | 0.011 | 1.83 | 41,42 |
| *GlcNAc* | R-HSA-8978868 | Fatty acid metabolism | 0.016 | 1.62 | 41,43 |
| *GlcNAc* | R-HSA-2022857 | Keratan sulfate degradation | 0.021 | 1.86 | 44 |
| *GlcNAc* | R-HSA-9612973 | Autophagy | 0.0063 | -1.68 | 6 |
| *GlcNAc* | R-HSA-6807878 | COPI-mediated anterograde transport | 0.011 | -1.73 | 45,46 |
| *GlcNAc* | R-HSA-5205647 | Mitophagy | 0.014 | -1.84 | 47 |
| *GlcNAc* | R-HSA-5205685 | PINK1 PRKN Mediated mitophagy | 0.02 | -1.85 | 47,48 |
| *GlcNAc* | R-HSA-9663891 | Selective autophagy | 0.025 | -1.66 | 6 |
| *GlcNAc* | R-HSA-1222556 | ROS and RNS production in phagocytes | 0.035 | -1.80 | 49 |
| *GlcNAc* | R-HSA-983168 | Antigen processing ubiquitination proteasome degradation | 0.043 | -1.46 | 6 |
| *Rem* | R-HSA-381119 | Unfolded protein response UPR | 0.0004 | 2.03 | 50 |
| *Rem* | R-HSA-381033 | Atf6 (atf6-alpha) activates chaperones | 0.0039 | 2.1 | 50 |
| *Rem* | R-HSA-5689880 | Ubiquitin specific processing proteases | 0.031 | 1.52 | 51 |
| *Rem* | R-HSA-5688426 | Deubiquitination | 0.046 | 1.44 | 51 |
| *Rem* | R-HSA-1630316 | Glycosaminoglycan metabolism | 0.0003 | -1.93 | 52 |
| *Rem* | R-HSA-1638091 | Heparan sulfate heparin hs gag metabolism | 0.0016 | -2.00 | 53 |
| *Rem* | R-HSA-1793185 | Chondroitin sulfate dermatan sulfate metabolism | 0.0093 | -1.80 | 54 |
| *Rem* | R-HSA-4085001 | Sialic acid metabolism | 0.014 | -1.82 | 55 |
| *Pli* | R-HSA-1852241 | Organelle biogenesis and maintenance | 0.045 | 1.42 | 46 |
| *Pli* | R-HSA-5619115 | Disorders of transmembrane transporters | 0.029 | 1.61 | 56 |
| *Pli* | R-HSA-9637687 | Suppression of phagosomal maturation | 0.0477 | 1.84 | 57 |
| *Pli* | R-HSA-425407 | SLC mediated transmembrane transport | 0.0024 | -1.73 | 56,58 |
| *Pli* | R-HSA-1483257 | Phospholipid metabolism | 0.0028 | -1.69 | 59 |
| *Pli* | R-HSA-9007101 | Rab regulation of trafficking | 0.0081 | -1.68 | 43 |
| *Pli* | R-HSA-2206281 | Mucopolysaccharidoses | 0.025 | -1.93 | 60 |
| *Pli* | R-HSA-1483206 | Glycerophospholipid biosynthesis | 0.027 | -1.56 | 59 |
| *Pli* | R-HSA-9840310 | Glycosphingolipid catabolism | 0.04 | -1.76 | 61 |
| *Pli* | R-HSA-917977 | Transferrin endocytosis and recycling | 0.046 | -1.81 | 62 |
| *SMLC* | R-HSA-191273 | Cholesterol biosynthesis | 0.0074 | 1.95 | 41 |
| *SMLC* | R-HSA-8957322 | Metabolism of steroids | 0.027 | 1.64 |  |
| *SMLC* | R-HSA-6811434 | COPI-dependent Golgi to ER retrograde traffic | 0.0040 | -1.86 | 45,46 |

Negatively enriched pathways are highlighted in light grey.

### Effects of drug candidates on lipofuscin content in CLN3 iNeurons and iAstrocytes

Since lipofuscin accumulation in brain cells is a pathological hallmark of Batten disease, we next examined whether drug candidates were able to modulate lipofuscin accumulation in individual cultures of CLN3 iPSC-derived iNeurons and iAstrocytes. The separate cultures were generated to investigate whether both cell types responded to drug treatment.

Phenotypical characterization of CLN3 iNeurons and iAstrocytes identified significantly reduced *CTSD* expression in CLN3 iNeurons and iAstrocytes and *LGALS3* in CLN3 iAstrocytes compared to CLN3-corrected cells (Supplementary Fig. 2a). There was also non-significantly reduction in *CLN3* expression in both CLN3 cell types, while no significant difference was identified for *LAMP1* expression (Supplementary Fig. 2a). As such, these results suggest that the aberrant lysosomal marker phenotype between CLN3 and CLN3-corrected cells was stronger in the mixed brain cell cultures, likely driven by both cell types.

Next, we tested the effects of candidate drugs on cell viability and lipofuscin content. The LDH release assay lipofuscin detection was performed following treatment with four drugs that elicited the highest number of gene expression changes and strongest pathway enrichment effects. The conditions were: 0.1 % DMSO (vehicle), GlcNAc 1 mM, Rem 10 µM, Pli 10 nM and SMLC 10 µM so that DMSO concentration within each conditions was the same. Interestingly, LDH release cytotoxicity assay demonstrated that in CLN3 iNeurons, Pli reduced cell death compared to UT, GlcNAc and SMLC while in contrast in iAstrocytes SMLC increased cell death compared to vehicle (Supplementary Fig. 2b). These results suggest that the mixed brain cultures had higher tolerance on drug treatments, where no differential effects on cytotoxicity were identified (Fig. 3a).

Lipofuscin was visualized via Sudan Black B (SBB) stain, which is considered a reliable stain for detecting lipofuscin in aging or senescent cells ^63,64^ as well as ceroid in Batten disease ^65^. Following drug exposure for 24 h, cells were fixed and stained with SBB along with a cell-specific marker (TUBB3 for iNeurons and S100B for iAstrocytes). The mean intensity for SBB in each image was then quantified and normalized to TUBB3 or S100B area (µm^2^). There were no significant differences in TUBB3 of S100B area between vehicle or drug treatment conditions in CLN3 iNeuron or iAstrocyte cultures, respectively, although S100B+ area was found to be significantly larger in UT CLN3 iAstrocytes compared to the other conditions (Supplementary Fig. 2c).

In CLN3 iNeurons, vehicle-only condition was found to significantly increase SBB signal intensity compared to UT (Supplementary Fig. 2d), suggesting potential sensitivity of iNeurons to DMSO treatment. Interestingly, GlcNAc treatment significantly (P < 0.01) reduced SBB intensity in the iNeurons, suggesting its ability to mitigate effects of DMSO on lipofuscin content within the cells (Fig. 6a). In CLN3 iAstrocytes, a non-significant elevation in SBB intensity was observed in the vehicle condition compared to UT (Supplementary Fig. 2d). Intriguingly, also in CLN3 iAstrocytes, GlcNAc (P < 0.01) treated cells exhibited significantly lower SBB intensity compared to vehicle, suggesting reduced lipofuscin within the cells (Fig. 6b). These results strongly indicate a protective role for GlcNAc in reducing lipofuscin burden in CLN3 brain cells.

**Figure 6.**
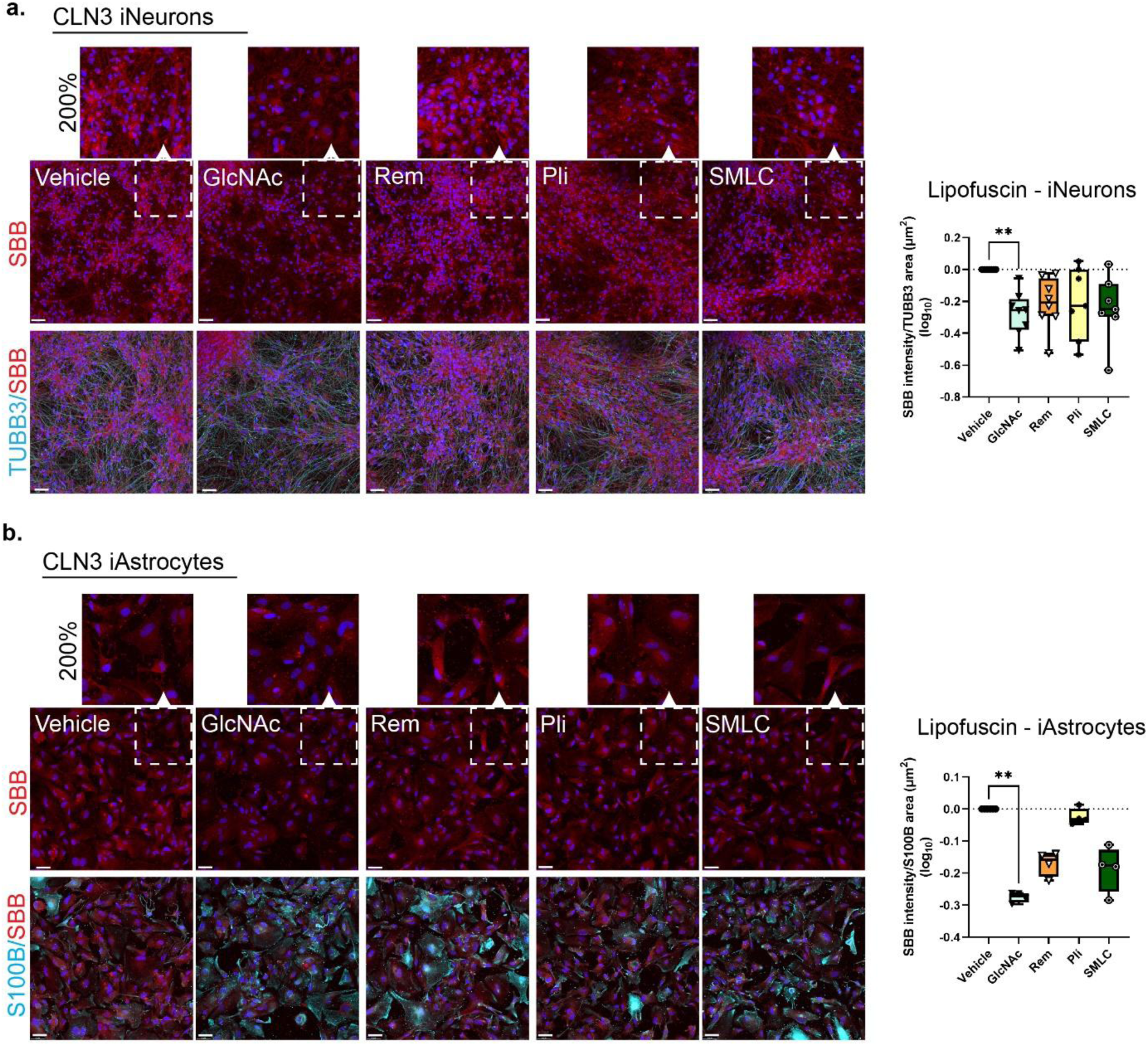
Effects of drug repurposing candidates on lipofuscin in CLN3 I Neurons and iAstrocytes. Fluorescent visualization of lipofuscin stained with Sudan Black B (SBB, red) (along with 200% zoomed-in sections) in GlcNAc, Rem, Pli and SMLC treated **a)** CLN3 iNeurons co-stained with TUBB3 (cyan) and **b)** CLN3 iAstrocytes co-stained with S100B (cyan) along with SBB mean intensity quantitation (presented as log_10_ of fold change of SBB/cell marker ratio). Statistical analysis was performed using Kruskal-Wallis test with Dunn’s multiple comparison correction and defined as ** P < 0.01.

## Discussion

Childhood dementia disorders are devastating neurodegenerative disorders for which there are limited treatment options and no cure. Therapies are available for less than 5 % of childhood dementia disorders, posing a highly unmet need ^66^. Batten disease is one of the most common causes of childhood dementia and is characterized by the accumulation of lipofuscin in brain cells. To accelerate the therapeutic discovery for Batten disease, we investigated the ability to employ computational drug repurposing combined with Batten disease-specific cell models to discover new therapeutic compounds.

Traditional drug development typically exceeds a decade and is marked by high failure rates. Drug repurposing provides a complementary, more efficient strategy by identifying new therapeutic uses for existing compounds, many of which already have established safety profiles from early-phase clinical trials ^67^. Importantly, drugs targeting genes with human genetic support are more likely to succeed in clinical development, underscoring the value of genetics-informed approaches ^68^. However, disease-associated genes rarely act in isolation. In Batten disease, we hypothesized that known pathogenic genes are likely to operate within tissue-specific molecular networks that influence disease onset and progression. We therefore applied a network biology framework ^69^ to map interactions between Batten disease causative genes and their functional partners, providing a systems-level view of disease mechanisms. This expanded network served as a molecular substrate for integrating drug-gene interaction data, enabling the identification of candidate compounds that may modulate disease-relevant pathways and offer opportunities for therapeutic repurposing.

Following drug repurposing analysis, we chose six drug candidates for further analysis, based on their effects on lysosomal enzymes and/or previous pre-clinical/clinical studies. Some of the selected compounds had clinical data reported on them, including GlcNAc for multiple sclerosis ^70^, Rem for COVID-19 ^25^, Pli for multiple myeloma ^71^ and Deoxy for Pompe disease (NCT04327973). In addition, SMLC is naturally present in garlic and onion with reported antioxidant effects ^72^ and MDL28170 has reported neuroprotective effects ^73^, thus they were considered as safe candidates for paediatric studies. The Lipinski’s rule of 5 filtering and low molecular weight supports BBB permeability, which is a bottle neck in brain-drug delivery ^74^. The overall safety of the candidate drugs was confirmed by observing minimal effects of on cell viability and inflammatory cytokine secretion. Whilst GlcNAc, Rem, Pli and Deoxy were found to significantly increase the secretion of the pro-inflammatory cytokine MCP1 (CCL2) secretion, the cytokine has been shown to increase neural stem cell self-renewal, proliferation and neuronal differentiation in a Niemann-Pick type C disease (another LSD) ^75^ and promote neuroblast migration after stroke ^76^. This suggests the roles of MCP1/CCL2 within the central nervous system (CNS) are more complex than what is understood.

As a limitation for disease-specific drug testing is the lack of appropriate disease-specific models, we utilized a patient-derived CLN3 brain cell model that consisted of a spontaneously differentiated mixture of neurons and astrocytes. This culture system provided a model for bulk brain tissue, which is a more reliable representation of the brain environment than simple brain cell mono-cultures, such as neurons. In particular, astrocytes play key roles in mediating neuron activity, neuroinflammation and regulating homeostasis ^77,78^, thus it is important to have these cells present in drug exposure studies to understand broader drug effects. Our approach of spontaneously differentiating CLN3 NPCs provides a fast and cost-effective drug testing platform for preliminary screening. For example, brain organoid models, which are considered important in brain drug testing, can be highly variable, time consuming and costly to generate ^79^. Importantly, our CLN3 brain cell model demonstrated a disease phenotype, confirmed via reduced gene expression of for *CLN3* and lysosomal markers *CTSD*, *LAMP1* and *LGAL3,* indicating their suitability for Batten disease-specific drug screening.

Although bulk RNA-seq revealed that none of the drug repurposing candidates significantly increased *CLN3* expression, GlcNAc was found to upregulate the *CLN5* gene, which encodes for the CLN5 protein, a soluble lysosomal glycoprotein ^80^. Defective CLN5 expression causes late-infantile Batten disease, although the role of CLN5 lysosomes is largely unknown ^80^. GlcNAc also upregulated lysosomal markers *LGALS3* and several cathepsins along with other genes associated with LSDs including *AGA* (aspartylglucosaminuria), *ARSA* (metachromatic leukodystrophy), *GALC* (Krabbe’s disease) and *HEXB* (Sandhoff disease), indicating lysosomal support. The downregulation of *CLN4* (*DNAJC5*) following GlcNAc and other drugs used in this study likely doesn’t have a detrimental effect, as CLN4 disease (also known as Kufs disease) is caused by mutant expression of *DNAJC5*, not by its deficiency ^81^. On the other hand, it would be important to confirm that the downregulation of CLN12 (*ATP13A2*), which is deficient in CLN12 disease (Kufor-Rakeb syndrome), does not have detrimental functional effects ^82^.

In terms of enriched pathways, GlcNAc and SMLC demonstrated similar effects, including the positive enrichment of ECM and collagen-related pathways. Collagen is the most abundant protein within the brain ECM and often considered a key scaffolding protein, however, in recent years its dynamic roles have become clearer, with the ability to bind other ECM proteins, cell receptors and ligands ^83^. For example, neurodegenerative diseases are associated with reduced collagen production, thus collagen-based therapeutics have been discussed as an option to promote neuronal regeneration in brain diseases ^83^. As such, collagen modifying and biosynthesis promoting compounds, could provide a new means to modulate collagen expression within the CNS. GlcNAc, Rem, Pli and SMLC treatments all shared negative enrichment in neurotransmitter release related pathways, which can indicate a reduction in neuronal activation, however, longer term exposures are required to investigate whether this effect persists past the 24 h timepoint.

Strongest support for further exploring the therapeutic use of GlcNAc in Batten disease comes from its effects on lipofuscin content in CLN3 iNeurons and iAstrocytes where GlcNAc significantly reduced lipofuscin/ceroid intensity within the cells. Considering that the effect of GlcNAc on lipofuscin was replicated across two separate cell types provides strong evidence for GlcNAc in protecting against cellular senescence and lipofuscin accumulation. The effects of GlcNAc on lipofuscin content could be driven by positive enrichment in the mitochondria fatty acid β-oxidation of unsaturated fats and fatty acid metabolism pathways, and negative enrichment in ROS production. Pathways associated with fatty acid oxidation and metabolism and oxidative stress are both linked to lipofuscin accumulation, as lipofuscin is largely composed of oxidised proteins and lipids ^84,85^. The therapeutic effects of GlcNAc are being investigated for other neurodegenerative diseases, including multiple sclerosis, with promising outcomes reported on GlcNAc reducing inflammation and neurodegeneration markers in patients ^70^. As a relatively low cost and safe compound, further investigation of GlcNAc in lysosomal storage disorders is warranted.

This study provides promising findings for a new candidate compound to be clinically investigated in Batten disease. Further expansion of the study findings to additional Batten disease models (including CLN5 Batten disease) will elucidate the potential benefits of GlcNAC on other forms of Batten disease and other lysosomal storage disorders.

Furthermore, longer exposure periods past the 24h used in the current study, will allow us to understand the long-term effects of GlcNAc treatment and identify whether some of the observed pathway changes are due to an acute response. Furthermore, single-cell transcriptomics and proteomics analysis will be important to identify the cell-type specific (neuron vs astrocyte) responses to GlcNAc.

## Methods

### Study design

The aim of this study was to apply a computational drug repurposing analysis to identify candidate drugs for Batten disease. Drug repurposing candidates were identified using a network propagation approach based on co-expression patterns with known Batten disease and lysosomal storage disorder genes. Drug candidates were prioritized based on clinical and experimental information, such as primary indications, clinical trial data, and chemical properties. An *in vitro* model of Batten disease (CLN3 type) was generated using spontaneous neural cultures generated from induced pluripotent stem cells (iPSCs) obtained from an individual with *CLN3* Batten disease ^28^. As control cell lines, isogenic iPSCs that were CRISPR/Cas9 corrected for the 966 bp deletion in the *CLN3* gene were used ^28^ along with HDFa an unrelated healthy control line ^29^. Differences in cell phenotype were assessed via immunofluorescence and real-time quantitative PCR (RT-qPCR). Bulk RNA-sequencing (RNA-seq) following 24 h drug exposure was used to identify differentially expressed genes and changes in cellular pathways. The effects of top drugs were further investigated on CLN3 iPSC-derived iNeuron and iAstrocyte lipofuscin content, the lysosomal storage material that accumulates in Batten disease.

### Computational drug repurposing analysis

Known genes implicated in Batten disease (Neuronal Ceroid Lipofuscinosis, NCL) were compiled from gene annotation databases (e.g., OMIM, ClinVar) and peer-reviewed literature. Genes including *CLN1* (PPT1), *CLN2* (TPP1), *CLN3*, and other *CLN* genes (*CLN4* to *CLN8* and *CLN10* to *CLN14*) with established pathogenic mutations linked to Batten disease were selected for subsequent analyses along with other childhood dementia genes (Supplementary Table 1.). Genome-wide co-expression networks specific to human brain tissues were generated using a data-driven Bayesian integration method, as previously described by Greene et al. ^86^. Briefly, publicly available transcriptomic data from brain tissues were integrated to construct brain-specific functional interaction networks. Co-expressed gene sets were identified based on high-confidence functional interactions (posterior probabilities ≥ 0.9) with known Batten disease genes. Network-based analyses, including functional enrichment (Gene Ontology terms, KEGG pathways), were conducted to characterize biological processes represented by these co-expressed gene sets. Genes co-expressed with known Batten disease causative genes were annotated for known drug-gene interactions using established pharmacogenomics and drug-target databases, including DrugBank ^87^ and the Drug-Gene Interaction Database (DGIdb) ^88^. Drug candidates annotated from co-expressed genes underwent additional prioritization by evaluating their molecular characteristics to predict BBB permeability using data in DrugBank. Specifically, we used Lipinski’s ‘rule of five’ to identify orally deliverable compounds based on the limits of four chemical properties (an orally active drug can have no more than one violation of the conditions): a molecular weight less than 500 Da, a calculated clogP (a measure of lipophilicity) less than 5, less than 5 hydrogen bond donors, and less than 10 hydrogen-bond acceptors ^15^. Drug candidates were selected based on prior clinical or promising *in vitro* effects.

### iPSC expansion and differentiation

The iPSC lines used in this study were generated and characterized and previously described ^28,29^. iPSC were expanded on human recombinant vitronectin (Thermo Fisher Scientific) in mTESRPlus medium (Stemcell technologies) and routinely characterized for the expression of pluripotency markers Nanog, OCT4 and SOX2 (Supplementary Figure 3a) and normal karyotype using the hPSC Genetic Analysis Kit (Stemcell technologies). For neural cell differentiation, iPSCs were first differentiated to neural progenitor cells (NPCs) using the STEMdiff^TM^ SMADi Neural Induction Kit (Stemcell technologies) with subsequent expansion of NPCs performed using STEMdiff^TM^ Neural Progenitor Medium. NPC identity was confirmed by the co-expression of Nestin and SOX2 (Supplementary Figure 3b). To generate spontaneously differentiated brain cell cultures, NPCs were seeded at density of 2 x 10^4^ cells/cm^2^ on 1:100 Matrigel (Corning) coated dishes in Neural Progenitor medium. The following day, culture medium was switched to Spontaneous Differentiation medium, consisting of BrainPhys^TM^ (Stemcell technologies), supplemented with B-27 and N-2 supplements (Thermo Fisher Scientific). Half medium changes were performed every 3 – 4 days and cells were differentiated for 30 – 33 days.

iAstrocyte differentiation was performed as previously described ^30^. Briefly, NPCs were seeded on Matrigel coated plates and culture medium switched to astrocyte differentiation medium (DMEM/F12 + GlutaMAX^TM^ + 1 % fetal bovine serum + N-2 supplement) and passaged when cells reached confluency. After approximately 40 days in culture, cells were seeded on plastic, which allowed cells to further mature. Cells were differentiated for 60 days prior to experiments and seven days before performing experiments, culture medium was supplemented with 10 ng/mL of bone morphogenetic protein 4 (BMP4) and ciliary neurotrophic factor (CNT) (Peprotech) to ensure further maturation.

iNeuron differentiation was initiated by seeding NPCs on poly-L-ornithine (20 µg/mL, Sigma-Aldrich) + laminin (10 µg/mL, Thermo Fisher Scientific) coating at a density of 2.5 x 10^4^ cells/cm^2^. Cells were allowed to attach overnight, after which culture medium was switched to neuron differentiation medium (BrainPhys^TM^ + N-2 + B-27 supplemented with 20 ng/ml BDNF and GDNF (Peprotech)). Half medium changes were performed every 3 – 4 days for 30 days.

### Drug Treatments

Six drugs were selected from the computational drug repurposing analysis and concentrations were selected based on a literature review: N-Acetyl-D-glucosamine (GlcNAc, Sigma) ^89-91^; Remdesivir (Rem, MedChemExpress) ^92,93^; Plitidepsin (Pli, MedChemExpress) ^94,95^; MDL-28170 (MDL, MedChemExpress) ^96,97^ and Deoxynojirimycin (Deoxy, MedChemExpress) ^98,99^, summarized in Table 1. Drugs were reconstituted in DMSO (Sigma) with DMSO-only conditions corresponding to the highest drug concentration included as a vehicle only condition. As such, two separate DMSO concentrations were used: 0.1 % and 1 %, which corresponded to the highest drug concentration for Rem, SMLC,

MDL and Deoxy as well as GlcNAc, respectively For drug treatments, medium was gently removed from the well and drug diluted in cell culture medium added to the cells. In addition to vehicle, untreated cells (UT), for which a normal medium change was performed, were included in each experiment. Drug treatments were incubated on the cells for 24 hours. After incubation, conditioned medium was collected for cytotoxicity and cytokine secretion assays and cell lysates were collected for RNA or cells were fixed for immunofluorescence.

### CyQUANT^TM^ LDH Cytotoxicity Assay

To investigate possible cytotoxic effects of drugs, lactate dehydrogenase (LDH) release was assessed using the CyQUANT^TM^ Cytotoxicity kit (Thermo Fisher Scientific). Briefly, following 24 h drug exposure, culture medium was collected from each treatment condition including a positive control for cell death, which had 1% of Triton-X 100 (Sigma-Aldrich) added to it 1 h prior to sample collection (100% cell death). Following sample collection, 25 μL of assay reaction mixture was added to each sample, and the contents were mixed by gentle tapping. The plate was incubated at room temperature for 30 min and protected from light. Following incubation, 25 μL of stop solution was added to each sample. Absorbance was measured at 490 nm and 680 nm using the Tecan infinite Control 2.0.10.0plate reader. To determine LDH activity, the 680 nm absorbance value was subtracted from the 490 nm absorbance value. To determine the percentage of dead cells, absorbance values were divided by the absorbance value of LDH positive control sample (100% cell death) and multiplied by a 100.

### RNA Extraction

For RNA collection, the culture media was removed from the wells and cells lysed with TRIzol^TM^ reagent (Thermo Fisher Scientific). Cells were scraped thoroughly off the culture plates and the lysates transferred into 1.5 mL Eppendorf tubes and stored at -80 °C until RNA extraction. For extraction, samples were kept on ice at the bench until fully thawed. A volume of 60 μL of chloroform (Sigma) was added to every 300 μL of Trizol. The mixture was vortexed vigorously for 15 seconds. Phase separation was achieved by centrifugation at 12,000 × g for 15 min at 4 °C, after which, the aqueous phase was carefully transferred to a new labeled tube. An equal volume of isopropanol (Sigma) was then added to sample tubes along with 0.5 μL glycogen (Thermo Fisher Scientific) to aid RNA precipitation. The samples were then incubated overnight at 4 °C. The next day, a second centrifugation was performed at 12,000 × g for 15 min at 4 °C. After centrifugation, the isopropanol was carefully removed using vacuum aspiration and 500 μL of 75% ethanol was added to each tube, and the tubes vortexed gently for 5 seconds. The samples were then centrifuged at 7,500 × g for 5 min 4 °C. The supernatant was removed as before, and another 500 μL of 75% ethanol was added. The tubes were vortexed until no residual liquid dripped out when the tubes were inverted. The RNA pellet was then resuspended in 20 μL of RNase-free injection-grade water. A water bath was prepared and maintained at 65 °C, in which the samples were placed for 10 minutes. After incubation, a quick spin (zip spin) was performed at maximum speed for 30 seconds. The RNA concentration and purity were measured using a Nanodrop spectrophotometer. Samples were stored at -80 °C until use.

### Bulk RNA-sequencing (RNA-seq)

For transcriptome analysis of drug- and vehicle-treated cells, bulk RNA-seq was performed at the QIMR Berghofer Next Generation Sequencing facility. For RNA-seq, for Pli, SMLC, MDL and Deoxy the highest concentration was used (10 nM or µM) and for GlcNAc 1 mM concentration was used to align with the same DMSO concentration as with the other drugs. For Rem the 5 µM concentration was selected due to some observations of the 10 µM concentration slightly increasing cell death and/or inflammation in cell models. Total RNA was extracted as described and quality determined using the Agilent TapeStation system.

The following samples underwent RNA-seq in triplicate (n = 3): vehicle (DMSO 0.01%), GlcNAc 1 mM, Rem 5 µM, Pli 10 nM, SMLC 10 µM, MDL 10 µM and Deoxy 10 µM. Library preparation was performed using Illumina TruSeq stranded mRNA Library prep kit and libraries were sequenced using the Illumina NextSeq 2000 P2 flow cell (single-end 100 cycles), resulting in an average of 19.92 million reads per sample (range 15.30 to 24.30 million). Sequence reads were trimmed for adaptor sequences using Cutadapt version 1.9 ^100^ and aligned using STAR version 2.7.10a ^101^ to the human GRCh38 assembly with the gene, transcript, and exon features of Ensembl release 110 gene model. All read-group aligned BAM files were merged using Samtools merge version 1.9 for each sample. Quality control metrics were computed using RNA-SeQC version 2.4.2 ^102^ and expression was estimated using RSEM version 1.2.30 ^37^, with the expected gene counts output used for downstream analysis. Differential expression analysis was performed using R version 4.5.1 and the quasi-likelihood pipeline from edgeR version 4.6.3 ^103-105^. Only protein-coding genes that passed the edgeR filterByExpr function were kept for further analysis. To compare each of the six drug-treated conditions to the vehicle condition, we used a design matrix without an intercept term, namely model.matrix(∼0 + Group), where Group was coded as a factor with the following levels: Vehicle, GlcNAc, Rem, Pli, SMLC, MDL, Deoxy. The edgeR glmQLFit function was used to fit a quasi-likelihood negative binomial generalized log-linear model to the read counts for each gene. The six contrasts of interest (GlcNAc – Vehicle, Rem – Vehicle, Pli – Vehicle, SMLC – Vehicle, MDL – Vehicle, Deoxy – Vehicle) were constructed using the makeContrasts function from limma version 3.64.1 ^106^. Using the edgeR glmQLFTest function, gene-wise empirical Bayes quasi-likelihood F-tests were conducted for each of the six contrasts. Differentially expressed genes (DEGs) between each of the six drug-treated conditions and vehicle were determined using a false discovery rate (FDR) < 0.05 calculated by the p.adjust function.

Gene set enrichment analysis (GSEA) ^107^ was performed on differential expression results for each experimental contrast to identify gene sets with coordinated, directional changes in gene expression. For each contrast, genes were ranked using a continuous, directional statistic derived from the edgeR quasi-likelihood F-test, defined as:

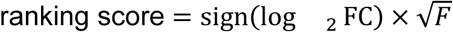

This score preserves both the direction of effect (with the sign function; up- or down-regulation) and statistical strength and approximates a t-like statistic suitable for pre-ranked GSEA.

Ranked gene lists were generated from the full set of tested genes without prior significance filtering and used as input for GSEA using WebGestaltR (v0.4.6) ^108^. Analyses were conducted for Homo sapiens using curated pathway databases, including Reactome ^109^.

Gene identifiers were mapped using HGNC gene symbols. Gene sets containing fewer than five genes were excluded from analysis. Enrichment significance was assessed using a running-sum statistic across the ranked gene list, with statistical significance estimated using 10,000 gene set permutations, and pathway-level enrichment summarized by normalized enrichment scores (NES). For each contrast, the top enriched pathways were reported based on rank-based significance criteria, consistent with the pre-ranked GSEA framework. All GSEA was performed independently for each experimental contrast and pathway database.

### cDNA synthesis and quantitative real-time (qRT-PCR)

The cDNA master mix was prepared for 100 ng of total RNA using the SensiFast^TM^ cDNA synthesis kit (Bioline) according to manufacturer’s instructions. To perform cDNA synthesis, the samples were placed in a thermal cycler, which was programmed as follows:

- 25 °C for 10 minutes for primer annealing
- 42 °C for 15 minutes for reverse transcription
- 85 °C for 5 minutes for enzyme inactivation
- Hold at 4 °C

After the run, samples were briefly centrifuged and diluted 1:10 with DNase/RNase-free H_2_O. For qRT-PCR run, the SensiFAST^TM^ SYBR® Lo-ROX kit (Bioline) was used with 2 µL cDNA sample and 400 nM forward and reverse primers of target genes (Supplementary Table 2) loaded into each well replicate. The qRT-PCR run was carried out using QuantStudio 5 (Thermo Fisher Scientific) using a two-step amplification protocol according to manufacturer’s instructions.

### Legendplex cytokine secretion assay

The concentration of cytokines and chemokines in cell culture condition medium was detected using the bead-based LEGENDplexTM Human Inflammation 13-plex panel (#740809) kit (BioLegend, CA, USA). The assay was performed as per the manufacturer’s instructions. Signal acquisition was performed using BD LSRFortessa 5 (BD Biosciences, CA, USA) flow cytometer using FACSDiva software and data were analysed using Qognit, a cloud-based software (BioLegend, USA).

### Immunofluorescence

For immunofluorescence, cells were cultured in a 24-well plate containing plastic coverslips or on plastic chamber slides (Sarstedt), or alternatively on black optical 96-well plates (Thermo Fisher Scientific), and fixed with 4% PFA for 15 min at room temperature. Permeabilization was performed with 0.3% Triton X-100 for 15 min, after which cells were blocked with 3% bovine serum albumin (BSA) in PBS for 2 h. Primary antibodies (Table X) were incubated overnight at 4 °C after which cells were rinsed 3 X with PBS. Secondary antibodies (Supplementary Table 2.) were then incubated for 2 h at room temperature. After incubation, secondary antibodies were removed and the wells were washed 2 X with PBS per wash after which Hoechst nuclear staining was performed for 5 min. After this, cells were washed two more times with PBS and mounted with ProLong^TM^ Gold antifade reagent (Thermo Fisher Scientific) or with Ibidi mounting medium. Confocal imaging of stained cells was performed with Leica STELLARIS confocal microscope.

### Sudan Black B lipofuscin stain

Sudan Black B stain (Sigma) was used to visualize lipofuscin as per a published protocol ^63^. Briefly, Sudan Black B was dissolved by adding 1.2 g of Sudan Black B to 80 mL of 70% ethanol in a glass bottle. The mixture was stirred overnight on a magnetic stirrer. Before use, the solution was filtered three times as follows: first through a 70 μm cell strainer, then through a 0.45 μm syringe filter, and finally through a 0.22 μm syringe filter. Sudan Black B stain was performed following secondary antibody stain and subsequent PBS rinsing. For this, the PBS solution was removed and the cells were incubated in 70% ethanol for 2 min. The ethanol was removed, and the cells were incubated in the freshly prepared Sudan Black B solution for 8 min. During staining, the plate was placed on a plate shaker set to 200 rpm. After staining, the Sudan Black B solution was removed and the cells were washed in distilled water for 5 min, with the plate again placed on the plate shaker at 200 rpm for the duration of the wash. The distilled water was removed, and a Hoechst nuclear stain was performed as described above. Sudan Black B staining was visualized using a Cy5 filter (far-red, 628/40 nm excitation; 692/40 nm emission) and quantified using ImageJ by calculating mean intensity. Mean intensity was divided by TUBB3+ or S100Bβ+ area determined by ImageJ.

### Statistical Analysis

Statistical analysis was performed using GraphPad Prism version 9.4.0. Comparisons between two groups were conducted using unpaired t-test. For comparisons involving more than two groups, ordinary one-way ANOVA was used, followed by appropriate posthoc tests when significant differences were found. ANOVA with Welch’s correction was used SDs were unequal. Kruskall-Wallis test was used when comparisons were made between groups when data was not normally distributed. A *P*-value of less than 0.05 was considered statistically significant.

## Acknowledgements

We would like to thank Batten Disease Support and Research Association Australia (BDSRA) for supporting this study. We would also like to thank Paul Collins and Tu Parsons at the QIMR Berghofer Next Generation Sequencing Facility for performing RNA-sequencing, Dean Basic (QIMR Berghofer) for performing the RNA-seq alignment and transcript quantitation and Sharon Hoyte (QIMR Berghofer) for assistance with the RNA-seq analysis. We would also like to thank Genome Informatics (QIMR Berghofer) for RNA-seq data management and computational support. We also acknowledge Oliver Looker (QIMR Berghofer Microscopy Facility and Spatial Cell Biology Facility) for assistance with immunofluorescence image analysis.

## Author contributions

L.E.O and Z.G designed the research. L.E.O designed and oversaw the cell experiments, analysed cell experimental data and wrote the manuscript. Z.G designed and performed the drug repurposing analysis, GSEA, generated figures and wrote sections of the manuscript. E.C, A.S.F and J.C performed the cell experiments, sample processing and downstream assays. E.C and A.S.F wrote sections for methods and Introduction. R.L.J performed RNAseq differential expression analysis and generated figures. D.A performed iPSC and NPC line characterization. S.C, A.L.C and A.W.H generated and provided the CLN3 and CLN3-corrected iPSC lines. E.M.D and A.R.W contributed to study design. All authors critically reviewed the manuscript and provided comments.

