## Supplementary information for "Computational drug repurposing identifies *N*-acetylglucosamine as a potential therapeutic compound for CLN3 Batten disease"

**Contents:**

Supplementary Figures 1 – 3

Supplementary Tables 1 – 4

**a.** Cytotoxicity

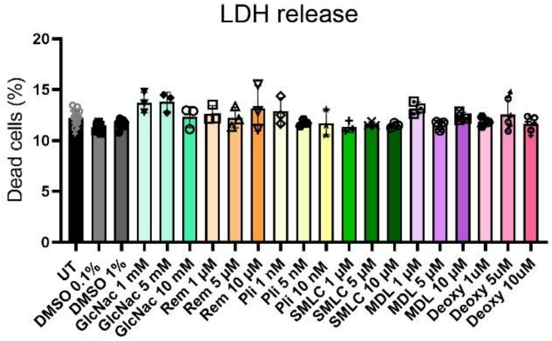


**b.** Neuroinflammation

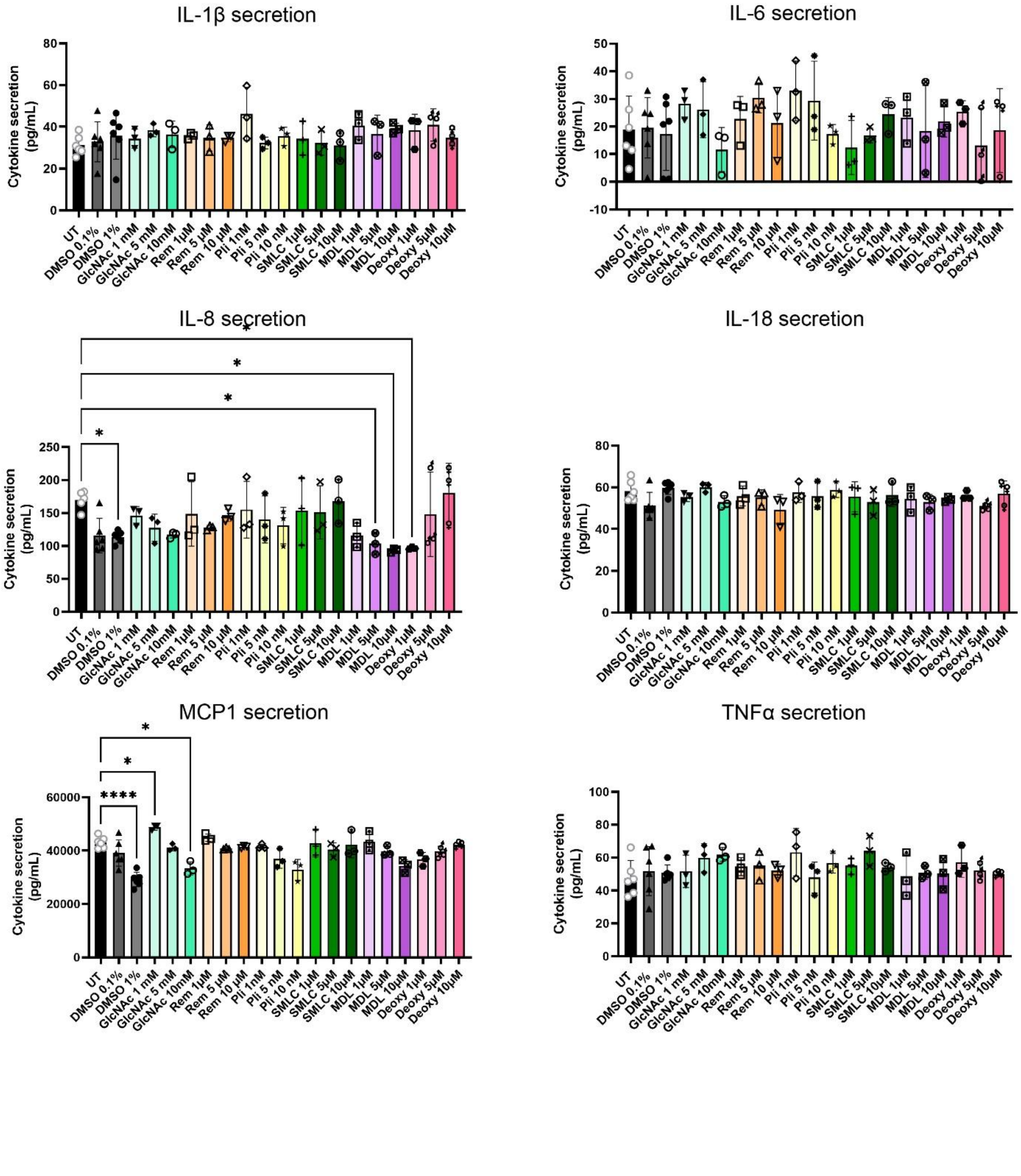

**Supplementary Figure 1. Effects of drug repurposing candidates on cytotoxicity and neuroinflammation compared to untreated condition. a)** Lactate dehydrogenase (LDH) release cytotoxicity assay following treatment with drug repurposing candidates in CLN3 brain cell model. **b)** Cytokine secretion in CLN3 brain cell model following treatment with drug repurposing candidates. Statistical analysis was performed with one-way ANOVA and defined as *$P < 0.05$, **$P < 0.01$ and ****$P < 0.0001$ (n = 3 independent replicates per treatment condition, only comparisons to untreated (UT) shown)).

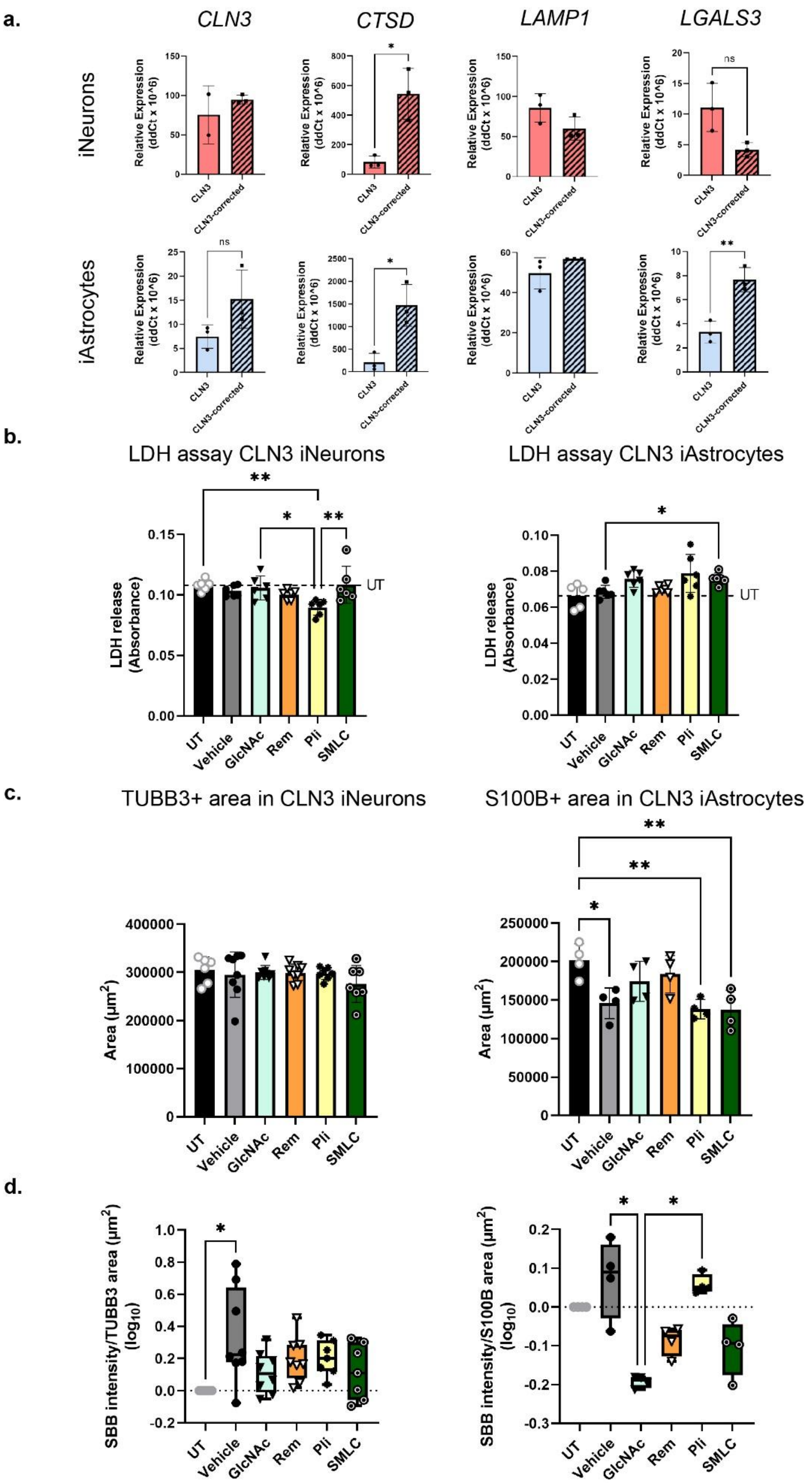
a.
CLN3
CTSD
LAMP1
LGALS3
iNeurons
iAstrocytes
Relative Expression (ddCt x 10^6)
CLN3
CLN3-corrected
ns
*
**
b.
LDH assay CLN3 iNeurons
LDH assay CLN3 iAstrocytes
LDH release (Absorbance)
UT
UT
Vehicle
GlcNAc
Rem
Pli
SMLC
c.
TUBB3+ area in CLN3 iNeurons
S100B+ area in CLN3 iAstrocytes
Area (µm²)
d.
SBB intensity/TUBB3 area (µm²) (log10)
SBB intensity/S100B area (µm²) (log10)

**Supplementary Figure 2. CLN3 iNeuron and iAstrocyte model characterization and lipofuscin quantitation. a)** Relative gene expression of lysosome markers CLN3 lysosomal/endosomal transmembrane protein (*CLN3*), cathepsin D (*CTSD*), lysosomal associated membrane protein (*LAMP1*) and galectin-3 (*LGALS3*) in CLN3 and CLN3-corrected iNeurons and iAstrocytes measured via qRT-PCR. **b)** Lactate dehydrogenase (LDH) release cytotoxicity assay following treatment with drug repurposing candidates in CLN3 iNeurons and iAstrocytes. **c)** Area quantitation ($\mu m^2$) of TUBB3 in CLN3 iNeurons and S100B in CLN3 iAstrocytes. **d)** SBB mean intensity quantitation (presented as $\log_{10}$ of fold change of SBB/cell marker ratio) in untreated (UT), vehicle and drug-treated iNeurons and iAstrocytes. Statistical analysis was performed with unpaired t-test for two groups or one-way ANOVA for more than two groups and defined as *$P < 0.05$, **$P < 0.01$ and ***$P < 0.001$ (n = 4 to 6 independent replicates per condition).

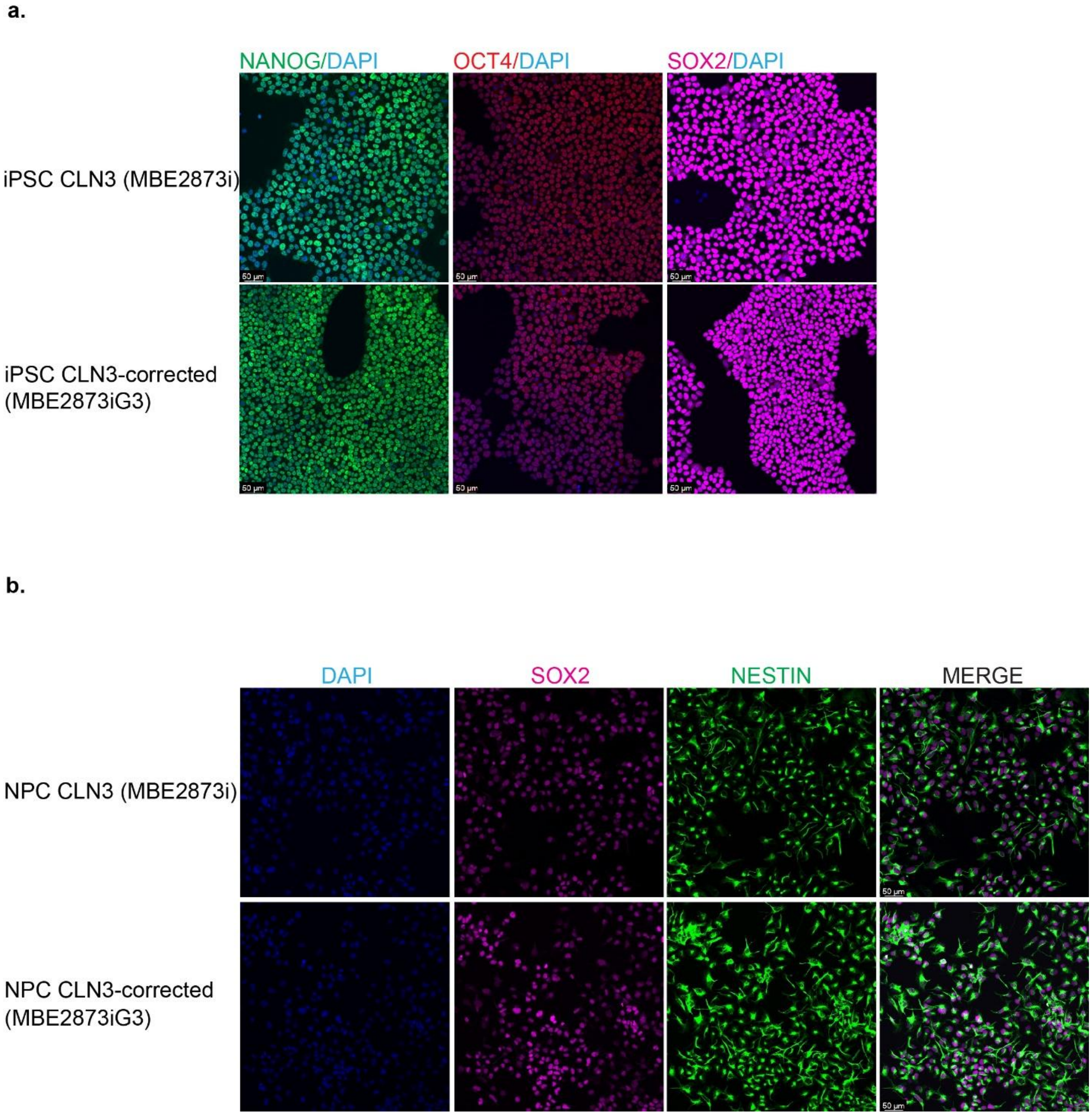


**Supplementary Figure 3. CLN3 (MBE2873i) and CLN3-corrected (MBE2873iG3) cell characterization. a)** Immunofluorescence of pluripotency markers Nanog (green), OCT4 (red) and SOX2 (magenta) in CLN3 and CLN3-corrected induced pluripotent stem cells (iPSCs). **b)** Immunofluorescence of co-expression of neural stem cell markers SOX2 (magenta) and Nestin (green) in CLN3 and CLN3-corrected iPSC-derived neural progenitor cells (NPCs).

**Supplementary Table 1. Input genes for drug repurposing analysis**

| Disease | Gene |
|---|---|
| Batten disease | *ATP13A2* |
| Batten disease | *CLN3* |
| Batten disease | *CLN5* |
| Batten disease | *CLN6* |
| Batten disease | *CLN8* |
| Batten disease | *CTSD* |
| Batten disease | *CTSF* |
| Batten disease | *DNAJC5* |
| Batten disease | *GRN* |
| Batten disease | *KCTD7* |
| Batten disease | *MFSD8* |
| Batten disease | *PPT1* |
| Batten disease | *TPP1* |
| Hurler Syndrome | *IDUA* |
| Sanfilippo disease | *HGSNAT* |
| Sanfilippo disease | *SGSH* |
| Sanfilippo disease | *NAGLU* |
| Sanfilippo disease | *GNS* |
| tay-sachs disease | *HEXA* |
| niemann-pick disease | *NPC1* |
| niemann-pick disease | *NPC2* |

**Supplementary Table 2. qRT-PCR primer sequences**

| Gene | Forward primer | Reverse primer | Accession Number |
|---|---|---|---|
| *CLN3* | TCTGTCTCTACGGCTGCTG | GGCCAAGAGGAGCCAACAAT | NM_000086.2 |
| *CTSD* | TGATTCAGGGCGAGTACATGA | ACACCTTGAGCGTGTAGTCC | NM_001909 |
| *DCX* | TATGCGCCGAAGCAAGTCTC | TACAGGTCCTTGTGCTTCCG | NM_000555.3 |

| | | | |
|---|---|---|---|
| *GFAP* | GAGGTTGAGAGGGACAATCTGG | GTGGCTTCATCTGCTTCCTGTC | NM_001242376.3 |
| *LAMP1* | GAAGGACAACACGACGGTGA | CATCCCGAACTGGAAGAGCA | NM_005561.4 |
| *LGALS3* | GCCAACGAGCGGAAAATGG | CAGGCCATCCTTGAGGGTTT | NM_002306.4 |
| *S100b* | TTCTGGAAGGGAGGGAGACA | CTCCTGCTCTTTGATTTCCTCT | NM_006272.3 |
| *TUBB3* | GGCCAAGTTCTGGGAAGTCAT | CTCGAGGCACGTACTTGTGA | NM_006086.4 |

**Supplementary Table 3. Immunofluorescence antibodies**

| **Antibody** | **Host species** | **Manufacturer** | **Catalogue number** | **Dilution** |
|---|---|---|---|---|
| AQP4 | Mouse | Abcam | Ab9512 | 1:100 |
| GFAP | Rabbit | DAKO | | 1:500 |
| MAP2 | Rabbit | Abcam | Ab281588 | 1:200 |
| Nanog | Rabbit | Abcam | Ab21624 | 1:100 |
| Nestin | Mouse | Abcam | Ab22035 | 1:200 |
| OCT4 | Rabbit | Abcam | Ab184665 | 1:100 |
| SOX2 | Rat | Invitrogen | 14-98110-82 | 1:100 |
| TUJ1/TUBB3 | Mouse | BioLegend | 801202 | 1:500 |
